# Reduced serotonergic transmission alters sensitivity to cost and reward via 5-HT_1A_ and 5-HT_1B_ receptors in monkeys

**DOI:** 10.1101/2023.02.23.529630

**Authors:** Yukiko Hori, Koki Mimura, Yuji Nagai, Yuki Hori, Katsushi Kumata, Ming-Rong Zhang, Tetsuya Suhara, Makoto Higuchi, Takafumi Minamimoto

## Abstract

Deficiency of the serotonin (5-HT) system is considered one of the core biological pathologies of depression and other psychiatric disorders whose key symptom is decreased motivation. Yet, the exact role of 5-HT in motivation remains controversial and elusive. Here, we pharmacologically manipulated the 5-HT system and quantified effects on motivation in terms of incentives and costs for goal-directed action in monkeys. Reversible inhibition of 5-HT synthesis increased refusal responses and reaction times in goal-directed task performance, indicating decreased motivation that could be separated into value-dependent and -independent components. To identify the receptor subtypes involved in these components, we systemically administered antagonists specific for four major 5-HT receptor subtypes: 5-HT_1A_, 5-HT_1B_, 5-HT_2A_, and 5-HT_4_. Positron emission tomography visualized the unique distribution of each subtype in limbic brain regions and determined the systemic antagonist dose that achieved approximately 30% occupancy. We found that blockade of 5-HT_1A_, but not other receptor subtypes, increased sensitivity to future workload and time-delay to reward, and decreased motivation in a value-independent manner. Moreover, blocking only 5-HT_1B_ receptors reduced the impact of incentive value on motivation. These results suggest that two distinct processes, mediated by 5-HT_1A_ and 5-HT_1B_ receptors, lead to reduced motivation in 5-HT system deficiency.

## Introduction

Generally, we make decisions about whether or not to engage in an action based on the trade-off between its benefits and costs – the expected value of the benefit (i.e., reward) has a positive influence, while the cost required to earn the expected reward (e.g., delay, risk, or effort) negatively influences the impact of the reward value (1-3). Deficiencies in the serotonin (5-HT) system are known to disturb this balance between value-related judgement and motivation. For example, because of its role as a major target of medication for depression and other psychiatric disorders affecting motivation, 5-HT deficiency is thought to be crucially involved in blunting or loss of pleasure in rewards, impulsivity to obtain rewards, and exhaustion of effort (4, 5). In rodent studies, 5-HT depletion increases impulsive choice (6-11). Attenuation of 5-HT transmission by specific receptor antagonists has been shown to increase impulsive choices and reduce the frequency of effortful behavior (12). In addition, pharmacological attenuation of dorsal raphe 5-HT neurons has been shown to impair waiting for long delayed rewards (13). Yet, despite this accumulating evidence, the specific mechanisms by which 5-HT contributes to motivation remain unclear, especially in primates.

Perhaps one of the challenges in understanding the role of 5-HT in motivation stems from the coexistence of two factors that modulate it: incentives and costs. Previous studies did not separately examine the incentives and costs for motivation, i.e., the behavioral effects of reducing the 5-HT system have never been quantitatively and independently examined in terms of incentive values and costs. Another complicating aspect of the 5-HT system is that there are fourteen receptor subtypes in the central nervous system (14) and their localization is heterogeneous. Therefore, it is important to characterize the role of each receptor in incentive- and cost-dependent modulations of motivation, as well as to localize them in the brain. In our previous study, we demonstrated dissociable roles of the two dopamine (DA) receptor subtypes in computation of the cost-benefit trade-off in motivation by combining positron emission tomography (PET) and pharmacological manipulation of DA receptors with quantitative measures of motivation in monkeys (15). Here, we extend this to the 5-HT system and set up experiments combining pharmacological manipulation with PET imaging to localize major 5-HT receptors and quantify drug actions in the brain.

In the present study, we aimed to elucidate the role of 5-HT neurotransmission in the motivation of goal-directed behavior. First, we manipulated the 5-HT system by repeated administration of para-chlorophenylalanine (pCPA), a reversible inhibitor of 5-HT synthesis, to macaque monkeys and examined its effects on goal-directed behavior. We further focused on four subtypes of 5-HT receptors (5-HTRs; specifically, 5-HT_1A_R, 5-HT_1B_R, 5-HT_2A_R, and 5-HT_4_R) that are abundant in limbic brain regions (16) by mapping their distribution, and manipulated 5-HT transmission by systemically administering antagonists of at doses predetermined by PET to achieve the same degree of receptor occupancy. Behavioral effects were assessed using two types of tasks designed to examine incentive impact and sensitivity to the costs (workload and delay) of obtaining a reward. Our results suggest that a reduction in 5-HT transmission leads to reduced motivation through two distinct processes: increased cost sensitivity via 5-HT_1A_R and reduced incentive motivation via 5-HT_1B_R.

## Results

### Effects of 5-HT depletion on incentive motivation

We first examined the effect of 5-HT depletion on incentive-driven motivation. We repeatedly injected pCPA (150 mg/kg, s.c.) for 2 days, which resulted in an approximately 50% decrease in 5-HT metabolites (5-hydroxyindole acetic acid, 5-HIAA) in the cerebrospinal fluid (CSF), while the concentration of DA remained at the same level (Figs 1A and 1B).

**Figure 1.**
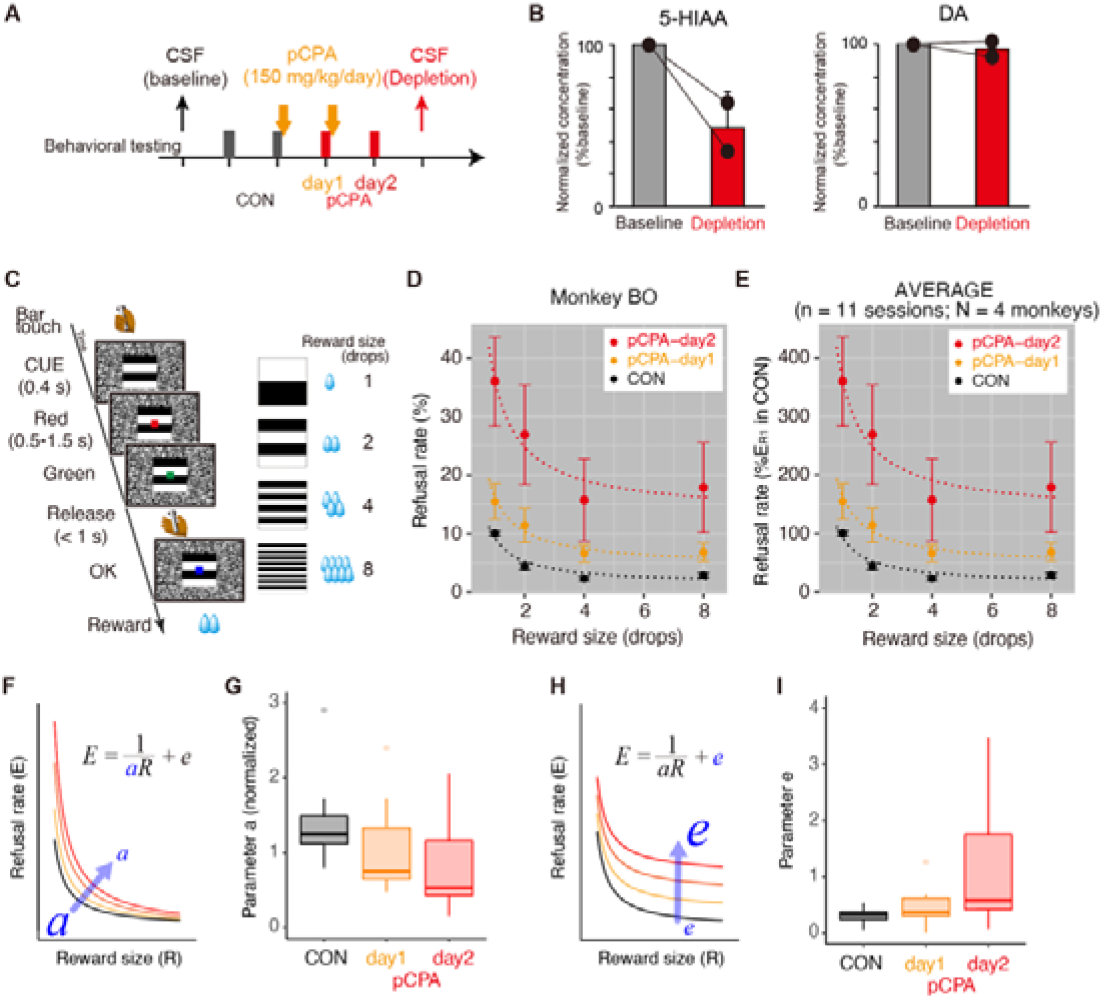
Effect of 5-HT depletion with pCPA on incentive motivation. (A) Schedule of pCPA treatment, behavioral testing, and cerebrospinal fluid (CSF) sampling. (B) Normalized concentration of 5-HT metabolite (5-hydroxyindole acetic acid, 5-HIAA) and DA before (baseline) and after pCPA treatment (depletion). (C) Reward-size task. Left: Sequence of events during one trial. Right: Relationship between visual cues and reward size. (D) Refusal rates (mean ± SEM) as a function of reward size for monkey BO. Dotted curves are the best-fit of inverse functions (Model #4, Table S2). (E) Normalized refusal rate (percent of maximum refusal rate in 1 drop trial in the control session; mean ± SEM) as a function of reward size for n = 11 sessions. (F) Schematic illustration of increase in refusal rate by reduction of incentive impact *a*. (G) Box plot of normalized incentive impact (*a*) for each treatment condition (n = 11 for each). Each value was normalized to the value of the control condition. (H) Schematic explaining the increase in refusal rate by *e*, independent of reward size. (I) Box plot of parameter *e* (normalized as ratio of maximum refusal rate in 1 drop trial in the control session; mean ± SEM) for each treatment condition (n = 11 for each).

We next examined the effects of 5-HT depletion on incentive motivation in four monkeys not used in the CSF study (Table S1). For this purpose, we used a reward-size task, the amount of reward was manipulated but the task requirements (i.e., the costs) were identical across trials (Fig 1C). For each trial of this task, the monkeys were required to release a bar when a visual target changed from red to green to receive a liquid reward. A visual cue indicated the amount of reward (1, 2, 4, or 8 drops) at the beginning of each trial. Based on previous studies using similar task, we measured the monkey’ s willingness to perform the task as a proxy for incentive motivation (2, 15). Thus, the frequency of refusal trials can be used as a behavioral measure of motivation. Furthermore, we show that the refusal rate (*E*) is inversely related to the reward size (*R*), which has been formulated with a single free parameter *a*,

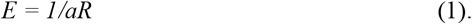

This inverse relationship was consistently observed in the control condition in all monkeys (e.g., CON in Figs 1D and 1E). With subsequent pCPA treatment, refusal rates increased regardless of reward size. For example, in monkey BO, the refusal rate became progressively higher for sessions following the first and second treatments, while differences that depend on reward size seem to be maintained (Fig 1D, pCPA-day 1 and 2). A reward-independent increase in refusal rate was consistently found in all monkeys tested, as shown in the average plot of normalized refusal rate (Fig 1E). We further quantified whether this increase was due to a reduction in the incentive impact of the reward or decrease in motivation independent of reward size. These factors can be captured by a decrease in parameter *a* of the inverse function and implementation of the intercept *e*, respectively (Fig 1F and 1H). To quantify the increase in refusal rate, we compared five models that considered these two factors as random effects: Model #1, random effect on *a*; Model #2, random effect on a fixed *e*; Model #3, independent random effects on both *a* and *e*; Model #4, random effects on *a* and *e*; and Model #5, random effect on *e* (see Table S2). Model #4 was selected as the best model with the lowest Akaike information criterion (AIC) value (Table S2), indicating that the increase in refusal rate was explained by simultaneous changes in parameters *a* and *e*. Our model-based analysis revealed that *a* significantly decreased in the pCPA-day 2 session compared with the control (one-way ANOVA, main effect of treatment, *F*_(2, 20)_ = 4.6, *p* = 0.023; post-hoc Tukey HSD, *p* = 0.025 for pCPA-day2 vs. CON; Fig 1G). Parameter *e* significantly increased in the pCPA-day 2 session compared with the control (main effect of treatment, *F*_(2, 20)_ = 4.1, *p* = 0.031; post-hoc Tukey HSD, *p* = 0.025 for pCPA-day2 vs. CON; Fig 1I). These results suggest that the 5-HT depletion-induced increased refusals can be explained by two components: one is a reduction of the incentive impact (*a*), and the other is a factor that appears orthogonal to the incentive value (increase in parameter *e*).

Given that the motivational value of reward decreases with increasing satiation (17), we further investigated how refusal increased along with satiation. In Figure 2A, the refusal rates of average data (n = 11) were replotted as a function of normalized cumulative reward (see Materials and Methods). As previously shown, overall refusal rates for each reward size increased as the normalized cumulative reward increased in the control condition. This satiation-dependent change in refusal rate was commonly observed among the three conditions, with the effect being stronger in the two post-treatment sessions (pCPA-day1 and -day2), reflecting a reduction of incentive impact. In post-treatment sessions, additional increases in refusal rates independent of satiation level were also pronounced; namely, refusal rates became higher even during the early phase of the session, presumably when the monkeys’ level of thirst drive should be high (Fig 2A). Indeed, fitting the data to a refusal rate model incorporating the satiation effect (Eq. 4) showed that refusal rate regardless of reward size or satiation level (*e*) was higher in post-treatment sessions (*e* = 3.3 and 6.6 for pCPA-day1 and -day2, respectively) compared with the control (*e* = 0.74). This suggests that 5-HT depletion blunts motivation in a manner partially independent of the motivational value, regardless of incentive or satiety.

**Figure 2.**
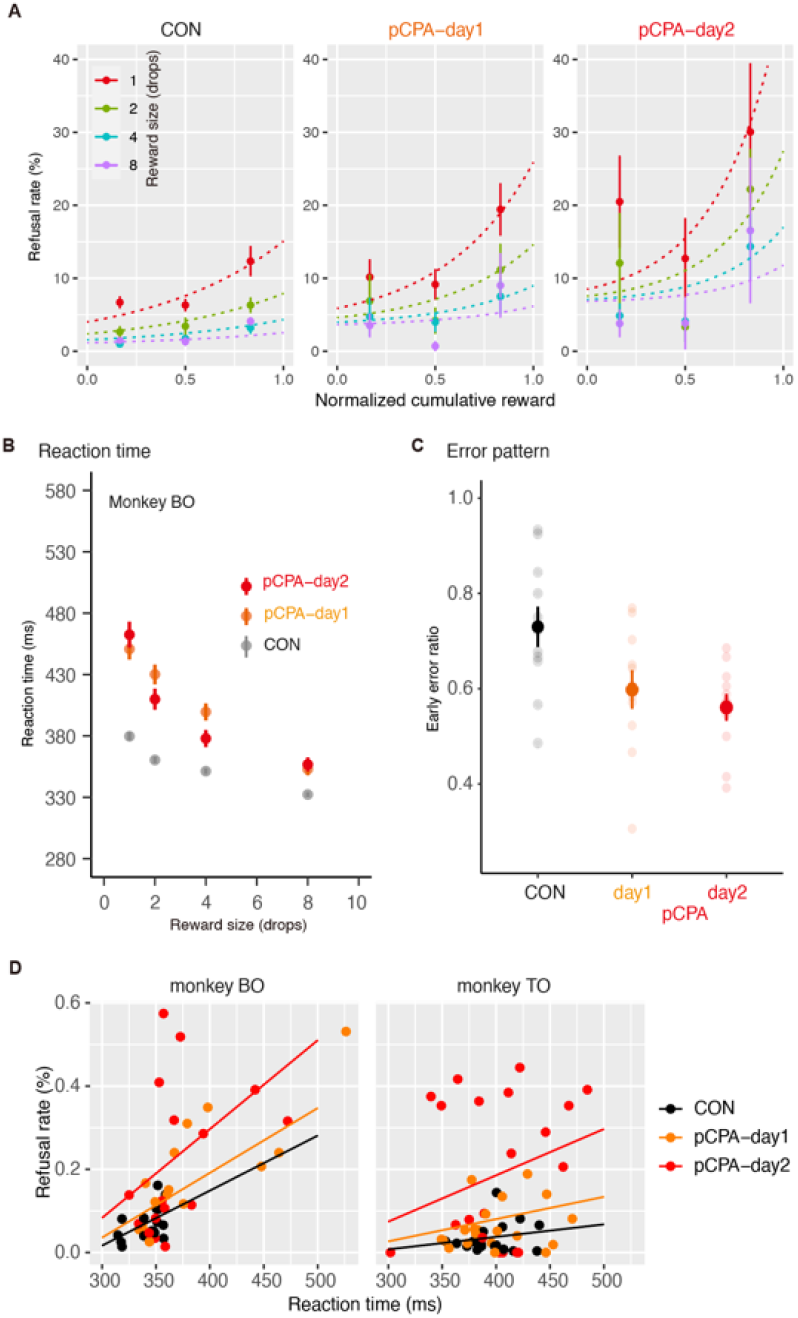
Effect of 5-HT depletion on impact of satiation on refusal, RT, error pattern, and RT-refusal relationship. (A) Refusal rate (mean ± SEM; n = 11 sessions) as a function of normalized cumulative reward for control and pCPA treatment conditions. Colors indicate reward size. Curves indicate best-fit model of Eq. 4. (B) Mean RT (mean ± SEM) as a function of reward size for control (CON) and pCPA treatment conditions in monkey BO. Colors indicate treatment condition. (C) Early release rate (mean ± SEM) for control and pCPA treatment conditions. (D) Relationship between refusal rate and mean reaction time for each reward size in session-by-session under pCPA treatment in monkeys BO and TO, respectively. Colors indicate treatment condition. Colored lines represent the best-fitting linear regression model to explain the data (Table S3).

We also examined the modulation of reaction time (RT) across trials as another behavioral measure of motivational process. Consistent with previous studies (18, 19), 5-HT depletion prolonged RTs (Fig 2B). Indeed, RTs were significantly increased with repeated administration [two-way ANOVA, main effect of treatment, *F*_(2, 110)_ = 7.1, *p* < 0.01; post-hoc Tukey’ s Honest Significant Difference (HSD), p < 0.01 for day1 vs. CON and day2 vs. CON] without significant interaction with reward size (main effect of reward size, *F*_(3, 110)_ = 6.0, *p* < 0.001 interaction, *F*_(6, 110)_ = 0.40, p = 0.88). pCPA administration also tended to decrease the proportion of early release (e.g., Fig 2C; one-way ANOVA, main effect of treatment, *F*_(2, 20)_ = 11.9, *p* < 0.001; post-hoc Tukey HSD, *p* < 0.005 for day1 vs. CON and day2 vs. CON). A simple explanation for these effects is that modulations in RT across conditions are caused by changes in motivation, such that the positive impact of reward on behavior affects both whether monkeys perform the action (refusal rate) and how quickly they will respond (RT). We reasoned that, if this were the case, the intersession variability in RT and refusal rate should be correlated. A session-by-session analysis revealed that although there was a linear relationship between refusal rates and RTs in each treatment condition, the slopes of the linear relationship became steeper as pCPA treatment was repeated (Fig 2D and Table S3). This finding suggests that additional factors beyond the normal motivational processes at work in decision-making and behavior may contribute to increased refusal.

### Effects of 5-HTR blockade on incentive motivation

Next, we sought to identify the receptor subtype(s) contributing to the value-independent decrease in motivation. We performed PET imaging with selective radioligands for 5-HT_1A_ ([^11^C]WAY100635), 5-HT_1B_ ([^11^C]AZ10419369), 5-HT_2A_ ([^18^F]altanserin), and 5-HT_4_ ([^11^C]SB207145), and quantified specific radioligand binding using a simplified reference tissue model (20) with cerebellum as the reference region. Consistent with previous human studies (16), different patterns of receptor distribution were observed for each subtype. For example, high levels of 5-HT_1A_R expression were observed in the limbic regions (including the anterior cingulate cortex, amygdala, and hippocampus), lateral prefrontal cortex, and posterior cingulate cortex, while expression was relatively sparce in the basal ganglia (Fig 3A). For 5-HT_1B_R, high expression was observed in the occipital cortex, ventral pallidum, and substantia nigra (Fig 3B). For 5-HT_2A_R, high expression was observed in the mediodorsal prefrontal and occipital cortex, while subcortical expression was minimal (Fig 3C). Binding of 5-HT_4_ was mainly observed in the striatum (Fig 3D).

**Figure 3.**
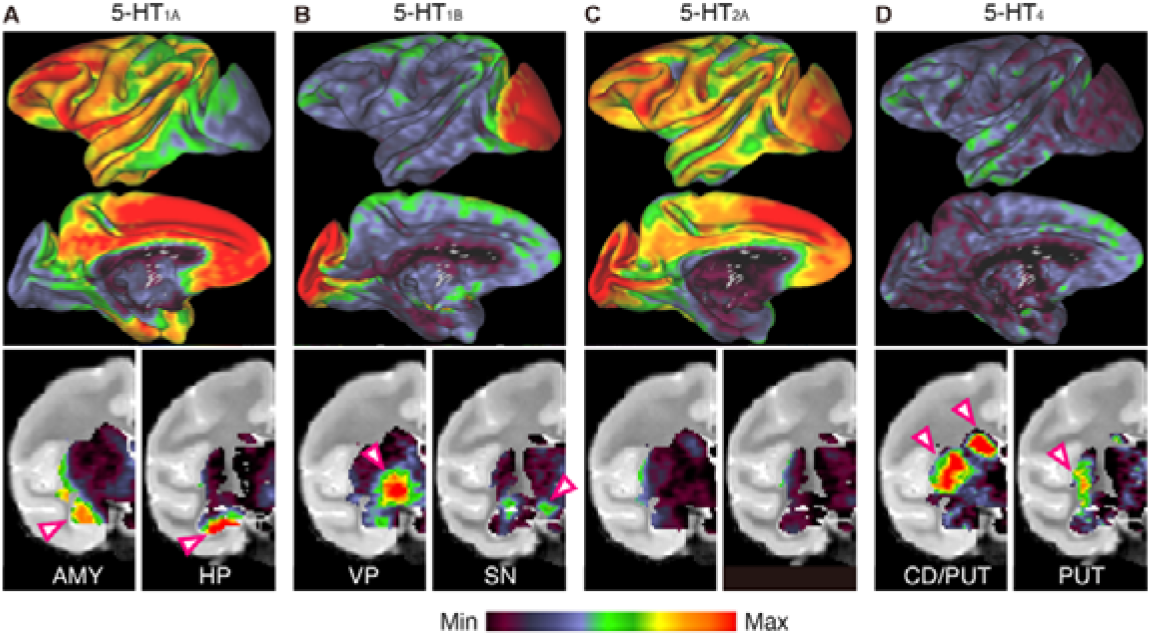
Heterogeneous distribution of 5-HTR subtypes as measured by PET. *Top*: Average density maps for four 5-HTR subtypes (A–D; 5-HT_1A_, 5-HT_1B_, 5-HT_2A_, and 5-HT_4_ receptors, respectively) on the inflated common macaque surface. Lateral (top) and medial (bottom) views of the left hemisphere are shown. *Bottom:* Subcortical distribution of four types of 5-HTRs shown on two representative coronal slices. Color scale represents 2^nd^–98^th^ percentile value of receptor density (i.e., radiotracer binding potential). AMY, amygdala; HP, hippocampus; VP, ventral pallidum; SN, substantia nigra; CD, caudate nucleus; PUT, putamen.

Selective antagonists are available for these four receptor subtypes, each of which is pharmacologically capable of attenuating or blocking 5-HT transmission. To compare the effect of receptor blockade between subtypes, it is critical to determine the appropriate drug dosage due to differences in pharmacological profiles such as receptor affinity and bioavailability. We previously used receptor occupancy as an objective measure for this purpose (15). To determine the appropriate antagonist dose of each receptor, we measured tracer binding at baseline and after systemic administration in four monkeys (Table S1), and inferred the degree of receptor blockade (i.e., receptor occupancy) (Fig S1). The dose of antagonist required to achieve approximately 30%–40% receptor occupancy was established, except for 5-HT_2A_, which achieved over 50% (Table 1).

**Table 1.** Antagonist dosage and occupancy for each 5-HT receptor subtype

| Receptor | PET ligand | Antagonist | Dose (mg/kg) | Occupancy |
| --- | --- | --- | --- | --- |
| 5-HT<sub>1A</sub> | [<sup>11</sup>C]WAY100635 | WAY100635 | 0.03 | 37% |
| 5-HT<sub>1B</sub> | [<sup>11</sup>C]AZ10419369 | GR55562 | 1 | 34% |
| 5-HT<sub>2A</sub> | [<sup>18</sup>F]altanserin | MDL100907 | 0.002 | 54% |
| 5-HT<sub>4</sub> | [<sup>11</sup>C]SB20715 | GR125487 | 1 | 32% |

We next evaluated the effects of blockade of each receptor subtype on performance in the reward size task in three monkeys, each of which was administered three of four different 5-HTR antagonists or vehicle as a control. Blockade of 5-HT_1A_R substantially increased the refusal rate (Fig 4A), while blockade of 5-HT_1B_R resulted in a moderate increase (Fig 4B). In contrast, blockade of 5-HT_2A_R or 5-HT_4_R did not induce any discernible change in refusal rate (Figs 4C and 4D). Model-based analysis revealed that blockade of 5-HT_1A_R consistently increased *e* in all monkeys (Model #3, Table S4), whereas blockade of 5-HT_1B_R, 5-HT_2A_R, and 5-HT_4_R induced an effect on *a* or individual influence (Model #6 or #10, Table S4). The best-fit models allowed us to extract parameter changes. As a result, blockade of 5-HT_1A_R increased the refusal rate regardless of reward size (*e*) by about 20% on average, while blockade of 5-HT_1B_R and 5-HT_4_R changed the incentive impact (*a*) to about 50% and 150%, respectively (Figs 4E and 4F). These results suggest that the two components of increased refusal rate observed upon 5-HT depletion can be reproduced separately by blocking two 5-HTRs: blocking 5-HT_1B_R reduces the incentive impact (*a*) and blocking 5-HT_1A_R increases a factor orthogonal to the incentive value (increased parameter *e*).

**Figure 4.**
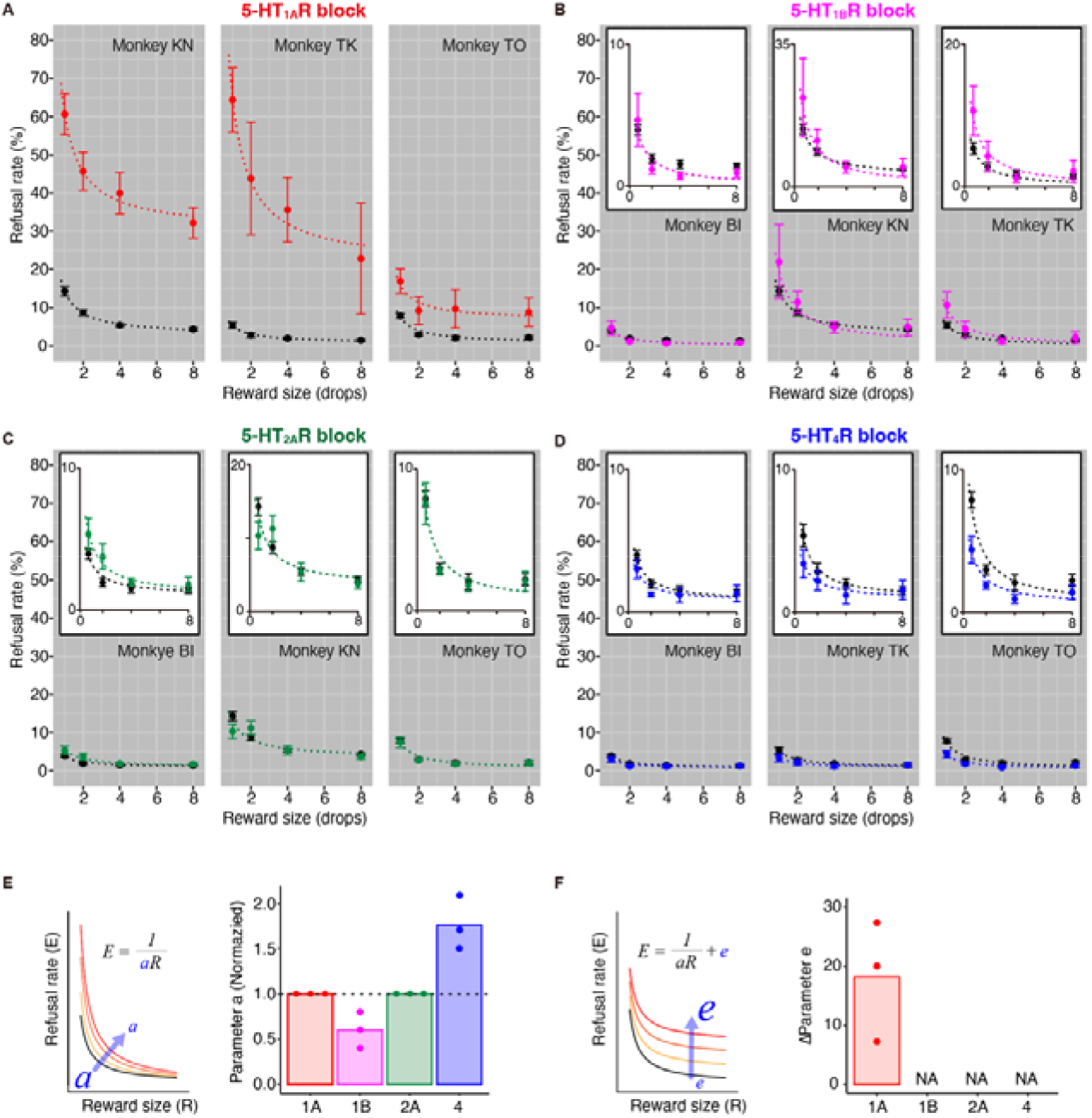
Effects of 5-HTR blockade on incentive motivation. (A–D) Refusal rates (mean ± SEM) as a function of reward size in three monkeys for blockade of 5-HT_1A_R, 5-HT_1B_R, 5-HT_2A_R, and 5-HT_4_R by systemic administration of specific antagonists. Dotted curves indicate the best fit of inverse functions summarized in Table S4. Rescaled plots are shown in insets in B–D. (E, F) Summary of parameter changes for *a* (incentive impact) and *e* (value-independent refusal), respectively. Dots and bars indicate individual data and the mean value, respectively. NA denotes that the parameter was not selected as a random effect in the best-fit model (S4 Table).

Blockade of 5-HT_1A_R prolonged RT independent of reward size (two-way ANOVA, main effect of treatment, *F*_(1, 214)_ = 366, *p* < 10^−16^; treatment × reward size, *F*_(1, 3)_ = 1.1, *p* = 0.33; Fig S2A). Blockade of any of the other 5-HTRs did not alter RT (*p* > 0.05; Figs. S2B and S2D). The error pattern did not change with any of the treatments (Fig S3). A session-by-session analysis revealed that the linear relationship between refusal rates and RTs was altered exclusively by 5-HT_1A_R blockade (Fig S4 and Table S5), again reproducing the behavioral changes observed with 5-HT depletion (Fig 2D). Thus, alteration of 5-HTR transmission via 5-H_1A_R appeared to mediate the deviation from normal value-based motivational processes seen with 5-HT depletion.

### Effects of 5-HTR blockade on cost-based motivation

Our results so far suggest that a reduction in 5-HT transmission via 5-HT_1A_R leads to a decrease in motivation independent of incentive values. One possible explanation for this phenomenon is that the effect may be mediated by factors orthogonal to incentive values in decision making, such as cost. Therefore, we investigated the effect of selective 5-HTR blockade on cost-based motivation. To accomplish this, we used a work/delay task (Fig 5A) that had the same basic features as the reward-size task, but instead manipulated two types of costs separately on each trial. In the work trials, the monkeys had to perform zero, one, or two additional instrumental trials to obtain a fixed amount of reward. In the delay trials, after the monkeys correctly performed an instrumental trial, a reward was delivered 0–7 seconds later. The number of trials or length of the delay was indicated by a visual cue presented throughout the trial. In the first trial after the reward, the visual cue indicated how much would have to be paid to get the next reward. Therefore, we assessed the monkeys’ performance on the first trials to evaluate the impact of expected cost on motivation and decision-making. We previously showed that monkeys exhibit linear relationships between refusal rate (*E*) and remaining cost (*CU*) for both work and delay trials, as follows:

**Figure 5.**
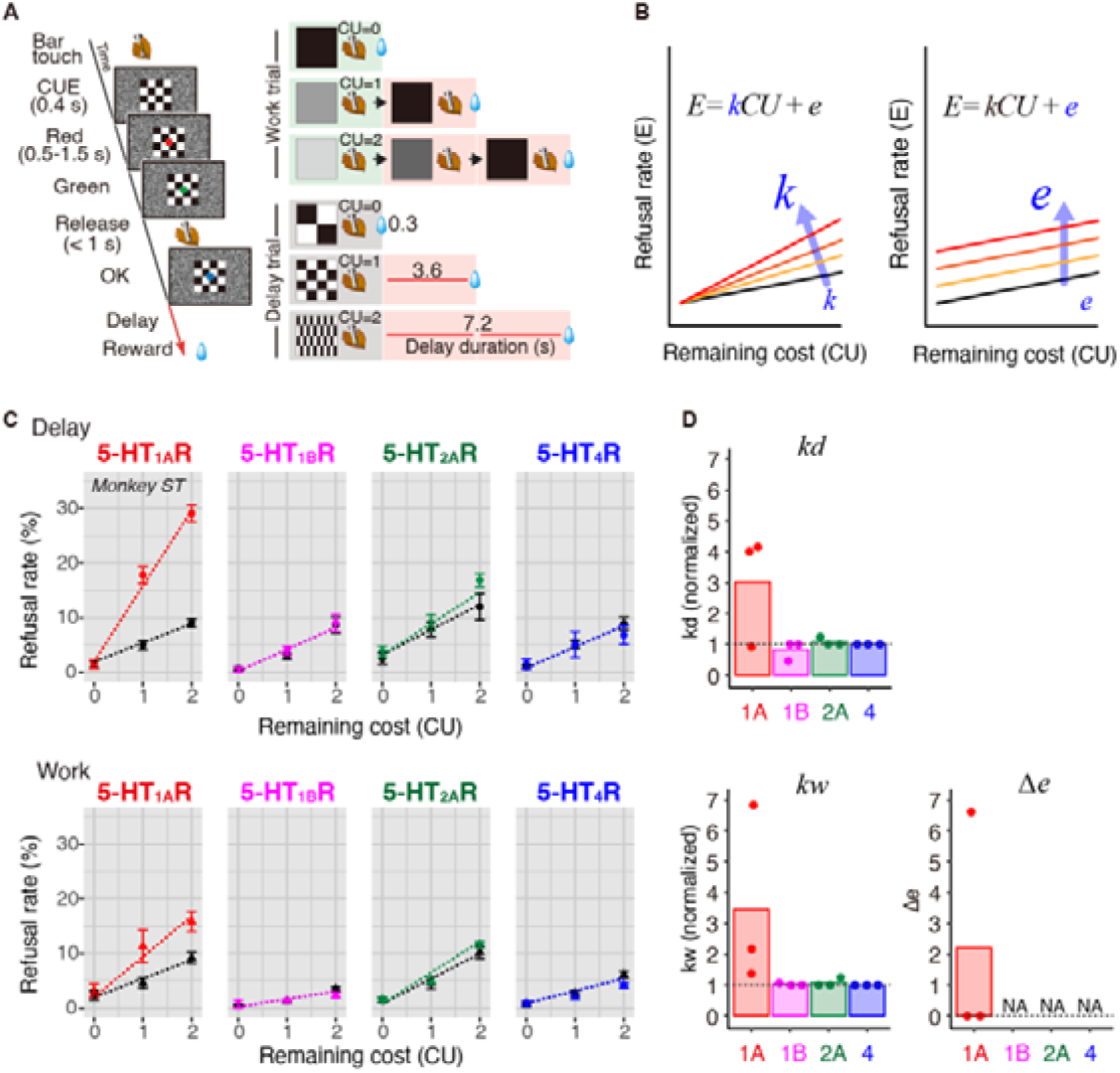
Effect of 5-HTR blockade on cost-based motivation. (A) The work/delay task. The sequence of events (left) and relationships between visual cues and trial timing in work trials (top three rows on the right) or delay duration in delay trials (bottom three rows on the left) are shown. CU denotes the remaining (arbitrary) cost unit to obtain a reward, i.e., either the remaining workload to perform the trial(s) or remaining delay periods. (B) Schematic of model for increases in refusal rate by increasing cost sensitivity (*k*) or increasing sensitivity to baseline cost (*e*). (C) Representative relationships between refusal rates (monkey ST; mean ± SEM) and remaining costs for delay (top) and work trials (bottom). Saline control (black line), 5-HT_1A_R, 5-HT_1B_R, 5-HT_2A_R, and 5-HT_4_R blockade (colored line) are shown from left to right, respectively. Colored and black dashed lines indicate the best-fit linear models described in S6 Table. (D) Comparison of the effects of 5-HTR blockade on the workload-discounting parameter (*k*_*w*_), delay-discounting parameter (*k*_*d*_), and sensitivity to baseline cost (*e)*.

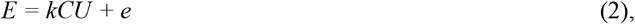

where *k* is a coefficient and *e* is an intercept (15) (Fig 5B). Extending the inference and formulation of the reward-size task (Eq. 1), this linear effect proposes that the reward value is hyperbolically discounted by the cost, where coefficient *k* corresponds to the discounting factors.

We tested three monkeys (Table S1) and measured their refusal rate to infer delay and workload discounting. We confirmed that the refusal rates of the control condition increased as the remaining cost increased (e.g., Fig 5C, control). Figure 5B illustrates two possible mechanisms by which 5-HTR blockade increases refusal rates. First, it may increase cost sensitivity (i.e., discounting factor, *k*), which appears as an increase in refusal rate relative to remaining cost (Fig 5B, left). The other is an increase in refusal irrespective to the degree of cost (i.e., increase *e*; Fig 5B, right). We again compared the effect of 5-HTR antagonism on task performance at the same level of receptor blockade (i.e., approximately 30% occupancy; Table 1). Linear mixed model (LMM) analysis tested the assumption that 5-HTR blockade independently increased delay and workload discounting without considering the random effect of treatment condition (e.g., Fig 5C and S6 Table; see Materials and Methods). We extracted the discounting factors and summarized their changes due to blockade in Figure 5D. Blockade of 5-HT_1A_R increased delay discounting in two of three monkeys and workload discounting in all three cases, whereas it only increased cost-independent refusal in one monkey (Fig 5D). Therefore, blockade of 5-HT_1B_R only decreased delay discounting in one of three cases. Blockade of other subtypes did not have a strong impact on discounting factors.

Blockade of 5-HT_1A_R prolonged RT independent of cost type (three-way ANOVA, main effect of treatment, *F*_(1, 358)_ = 92.2, *p* < 10^−16^; treatment × remaining cost, *F*_(1, 2)_ = 6.1, *p* = 0.002; treatment × trial type, *F*_(1, 1)_ = 3.6, *p* = 0.057; Fig S5A), except for in one monkey (ST). In contrast, blockade of the other subtypes did not prolong RT (*p* > 0.05; Figs S5B–D). None of the treatments changed the error pattern (*p* > 0.05; Fig S6).

## Discussion

5-HT deficiency has been shown to decrease motivation and alter cost sensitivity in humans. Studies in rodents have provided insights into roles of 5-HT signaling in various aspects of motivational behavior, but a gap remains between these basic findings and behavioral disorders in humans. Using macaque monkeys, the present study aimed to fill this gap and identify specific roles of 5-HTR subtypes in reduced motivation. We found that a reduction in 5-HT transmission leads to reduced motivation through two distinct processes: increased cost sensitivity via 5-HT_1A_R and reduced incentive motivation via 5-HT_1B_R. Our findings shed light on the mechanistic role of 5-HT in behavioral regulation and its relevance to depression medications.

### Two factors decreasing motivation of goal-directed behavior

Previous behavioral pharmacological studies demonstrated that 5-HT depletion slows action and/or reduces the likelihood of engaging in behavior (18, 19, 21), suggesting decreased motivation. However, these previous studies did not examine the effect of 5-HT depletion on incentive motivation in terms of multiple incentive conditions. Thus, data describing the quantitative relationship among 5-HT, reward, and motivation are unavailable. In the present study, we formulated and quantified the relationship using a behavioral paradigm (*reward-size task*). Using the same paradigm, we previously demonstrated that DA receptor blockade (either D1-or D2-like receptors) increased the rate of refusal, which was explained by a decrease in the impact of reward on motivation (reduced parameter *a*) (15). In the current study, 5-HT depletion also increased the refusal rate; however, this was partly explained by a reduction of the incentive impact, as well as a factor orthogonal to the incentive value (increased parameter *e*). In addition, refusal rates increased constantly regardless of the satiation level, suggesting a decrease in motivation independent of incentive value. The two factors of motivation reduction, namely, incentive-dependent and -independent factors, emerged specifically and exclusively after 5-HT_1B_R and 5-HT_1A_R antagonism, respectively. These behavioral effects could be directly compared among four major 5-HTR subtypes because we controlled their doses to achieve the same level of occupancy using PET. Thus, our findings suggest two independent mechanisms for 5-HT depletion-induced motivational decrements that are distinctively mediated by two receptor subtypes (5-HT_1A_R and 5-HT_1B_R) distributed in different brain regions, as shown by PET imaging. As discussed below, our results highlight a new view of the 5-HT system at the receptor level with respect to motivational decrements.

### Reduced 5-HT transmission via 5-HT_1A_R leads to decreased motivation due to overestimation of future costs

One of the key findings of this study was that blockade of 5-HT_1A_R leads to a deviation from the normal incentive-motivational process (or incentive-independent decrease in motivation), partially analogous to that seen during 5-HT depletion. This includes an increase in refusal rates and RT independent of reward size, and a change in the refusal-RT relationship. This incentive-independent change may be related to an overestimation of impeding cost; when we manipulated costs (forthcoming workload or delay duration to reward) independently of incentive value, the impact of cost on motivation specifically increased following 5-HT_1A_R blockade (Fig 5). These observations led us to conclude that reduced 5-HT transmission via 5-HT_1A_R results in increased cost sensitivity and decreased motivation.

5-HT has been implicated in the control of temporal discounting, which is the part of the cost-benefit evaluation process that regulates motivation. In humans, a decrease in 5-HT caused by tryptophan depletion has been shown to increase the rate of delayed reward discounting (9, 22). Conversely, 5-HT reuptake inhibitors (SSRIs, presumably by upregulating 5-HT) decrease delay discounting (23). Similarly, 5-HT depletion induced by pCPA administration tended to cause rats to choose immediate small rewards over long delay large rewards (7), whereas administration of an SSRI was associated with a decrease in impulsive choice (6). In addition to 5-HT depletion, reduced 5-HT transmission mediated by 5-HT_1A_R results in increased cost sensitivity to delayed reward; specifically, administration of an 5-HT_1A_R antagonist (WAY100635) tended to cause rats to choose immediate small rewards over prolonged delayed large rewards (6). In monkeys, 5-HT_1A_R blockade decreased correct responding as the delay to reward increased (12). However, other studies suggest mixed results, with relatively weak or no effect of 5-HT_1A_R antagonism on delayed discounting (24-26). This may be because these studies confounded incentive and cost factors in each option.

In contrast to delay sensitivity, the involvement of 5-HT in valuation of other types of costs, such as effort, is relatively controversial. A rodent study of cost-benefit trade-offs showed that SSRIs reduced effort expenditure (27), whereas 5-HT depletion with pCPA had no effect on the tendency of rats to make cost-based choices (7, 28). However, a human study showed that subjects taking SSRIs produced more effort for monetary incentives, which was mediated by a reduction in the cost of effort without changing in the weight of incentives (5).

As our PET data show (Fig 3), 5-HT_1A_Rs are predominantly expressed in the limbic system, such as the medial prefrontal cortex (mPFC), amygdala, and hippocampus, which is in good agreement with human data (16). The binding potential of 5-HT_1A_R, including these regions, is diminished in patients with major depressive disorder (29, 30) and monkeys exhibiting depression-like behavior (31), suggesting that alteration of 5-HT transmission is a pathophysiology associated with depression and probably its behavioral phenotype, including effortful information processing. However, additional studies are needed to identify the brain circuity contributing to increased cost sensitivity via 5-HT_1A_R. In the current study, limited or no involvement in increase of cost-sensitivity was found for other receptor subtypes. However, several previous rodent studies suggest a contribution of 5-HT_2A_R in the mPFC to impulsivity (32-34).

### Reduced 5-HT transmission via 5-HT_1B_R leads to decreased motivation in a value-dependent manner

Another component of motivation decline, an incentive-dependent one, was found to be reproduced by 5-HT_1B_R blockade. Importantly, this effect was independent of cost sensitivity and, thus, mediated by reward valuation or incentive function. Similar value-dependent reductions in motivation were observed following DA receptor antagonism with either D_1_R or D_2_R blockade (15). Given the mutual interactions between 5-HT and DA systems (35, 36), the behavioral changes observed here may be mediated, at least in part, by DA. 5-HT_1B_R is abundant in the basal ganglia, particularly in the output structures, globus pallidus, and substantia nigra (Fig 3B). Moreover, activation of 5-HT_1B_R increases DA release in the mesocorticolimbic system in rats (37, 38).

Additionally, several lines of evidence suggest a contribution of 5-HT_1B_R to incentive motivation. Blockade of 5-HT_1B_R attenuates cocaine-seeking, but not natural food-intake (39), suggesting 5-HT transmission via 5-H_1B_R is related to outcome-based motivational control. Our findings extend the current view of 5-HT_1B_R function, derived primarily from rodent studies, to primates and emphasize that its contribution to motivation is reward-related rather than cost-related (40). This has important implications for understanding how 5-HT regulates motivational behavior and its relationship to the pathophysiology and pharmacotherapy of depression and other disorders. A previous study suggested that upregulation of postsynaptic 5-HT_1B_R in the nucleus accumbens and ventral pallidum may be involved in the antidepressant effect of ketamine (41). Alteration of 5-HT_1B_R binding has also been reported in some psychiatric disorders, such as post-traumatic stress disorder (PTSD), alcohol dependence, pathological gambling, and drug abuse (42-46). It will be important to conduct future studies focusing on the relationship between altered 5-HT_1B_R function and reward sensitivity in humans.

### Limitation of this study

Because of the use of systemic antagonist administration, the current study could not determine which brain area(s) is responsible for antagonist-induced changes in incentive and cost sensitivity. While our findings, particularly the differential localization of receptor subtypes (Fig 3), support the idea that different limbic structures are involved in incentive and cost sensitivity, further research (such as local infusion of 5-HTR antagonists) is needed to identify key loci and generalize our findings to elucidate the circuit and molecular mechanisms of motivation.

Second, because our receptor blockade was partial and incomplete, the motivational involvement of other receptor subtypes may have been overlooked. For example, previous studies reported the involvement of 5-HT_2A_R in temporal discounting; specifically, systemic administration of the mixed 5-HT_2A_R agonist DOI dose-dependently impaired the ability to wait, whereas the 5-HT_2A_R antagonist ketanserin blocked the impulsivity effect of DOI (47). In the present study, relatively high blockade of 5-HT_2A_R (50% occupancy) did not alter incentive motivation or cost sensitivity.

However, this does not rule out the possible involvement of 5-HT_2A_R in these functions, as has been previously suggested.

### Conclusion

The present study demonstrates a differential contribution of receptor subtypes to reduced motivation caused by decreased 5-HT transmission: 5-HT_1A_R mediates increased cost sensitivity, whereas 5-HT_1B_R mediates reduced incentive sensitivity. Because these two receptor subtypes are differentially distributed in limbic brain areas, functional dissociations may reflect different circuit mechanisms of 5-HT on valuation processes that regulate motivation for action. Taken together, our findings add an important aspect to current knowledge of the role of 5-HT signaling in motivation based on cost-benefit trade-offs, thus providing an advanced framework for understanding the pathophysiology and medication of psychiatric disorders.

## Materials and Methods

### Ethics statement

All experimental procedures involving animals were conducted in accordance with the Guide for the Care and Use of Nonhuman Primates in Neuroscience Research (The Japan Neuroscience Society; http://www.jnss.org/en/animal_primates)” www.jnss.org/en/animal_primates) and approved by the Animal Ethics Committee of the National Institutes for Quantum Science and Technology (#09-1035).

### Subjects

A total of 16 adult male macaque monkeys (14 Rhesus and 2 Japanese; 4.5–7.9 kg; aged 4–12 years at the start of the experiment; see Table S1 for a summary of subjects) were used in this study. Food was available ad libitum, and motivation was controlled by restricting access to fluid to experimental sessions, in which water was provided as a reward for task performance. Animals received water supplementation whenever necessary (e.g., when they were unable to obtain sufficient water through experimentation) and had free access to water whenever testing was interrupted for more than 1 week. For environmental enrichment, play objects and/or small foods (fruits, nuts, and vegetables) were provided daily in the home cage.

### Drug treatment

For 5-HT depletion, monkeys were injected intraperitoneally with pCPA solution (C3635, Sigma-Aldrich, 0.9% in saline) at a dose of 150 mg/kg/day for two consecutive days. For 5-HTR blockade, the following 5-HTR antagonists were used: WAY100635 (W108, Sigma-Aldrich; for 5-HT_1A_R), GR55562 (cat# 1054, Tocris; for 5-HT_1B_R), MDL100907 (M3324, Sigma-Aldrich; for 5-HT_2A_R) and GR125487 (cat# 1658, Tocris; for 5-HT_4_R). WAY100635, GR55562, and GR125487 were dissolved in 0.9% saline, while MDL100907 was suspended in a drop of hydrochloric acid and the final volume was adjusted with saline. The dose for each antagonist is listed in Table 1. Monkeys were pretreated with either antagonist intramuscularly 15 min prior to the start of the behavioral test or PET scan. For behavioral testing, saline was injected as a control using the same procedure as for drug treatment. The volume of administered was set at 1 mL for all experiments.

5-HTR blockade was performed four times per individual antagonist in the behavioral test. Each vehicle or antagonist was administered once per week, with days of the week of administration counterbalanced. After completion of an antagonist test, the animals’ baseline task performance and condition were assessed, including daily activity in the home cage, body weight, and water and food consumption. If there were no abnormalities, the next sequence of behavioral tests was started with a different antagonist. The order of treatment of the four antagonists was counterbalanced for each animal.

### Measuring 5-HT metabolites and DA in CSF

Acute CSF samples were collected by lumbar puncture with a 23-gauge needle under ketamine-xylazine anesthesia. CSF was collected twice (0.5 µL/sample per monkey): once a week before pCPA administration and again the day after two consecutive days of pCPA administration. The supernatant was passed through a 0.22-μm filter (Millex-GV, Merck, Germany), centrifuged at 2,190 × g for 20 min, and aliquoted into another microcentrifuge tube. For analysis of 5-HIAA and DA, high-performance liquid chromatography (HTEC-500, EICOM Co., Japan) was used with a monoamine separation column (SC-5ODS, EICOM). CSF data were recorded and analyzed using Power Chrom software (version 2.5, eDAQ, USA). The mobile phase was a mixture of 0.1 M phosphate buffer (pH 6.0) and methanol at a ratio of 83:17, including EDTA-2Na (5 mg/L). The amount of monoamine was calculated quantitatively as an absolute amount relative to the monoamine standard solution (MA11-STD, EICOM).

### PET procedure and occupancy measurement

Four monkeys were used for PET measurements, which were performed with four PET ligands: [^11^C]WAY100635 (for 5-HT_1A_R), [^11^C]AZ10419369 (for 5-HT_1B_R), [^18^F]altanserin (for 5-HT_2A_R), and [^11^C]SB207145 (for 5-HT_4_R). PET scans were performed using an SHR-7700 PET scanner (Hamamatsu Photonics Inc., Japan) or microPET Focus220 scanner (Siemens Medical Solutions USA) on monkeys under conscious conditions and seated in a chair. The injected radioactivity and its molar activities at the time of injection were 88.4∼360.4 MBq and 14.8∼296.3 GBq/µmol, respectively. After transmission scans for attenuation correction, a dynamic emission scan was performed for 90 min, except for scans with [^11^C]SB207145 (120 min). Ligands were injected as a single bolus via the crural vein at the start of the scan. All emission data were reconstructed by means of filtered back projection using a Colsher or Hanning filter. Tissue radioactive concentrations were obtained from volumes of interest (VOIs) placed on the global cortex, visual cortex, and cerebellum (as a reference region). Each VOI was defined on individual structural magnetic resonance (MR) images (EXCELART/VG Pianissimo at 1.0 Tesla, Toshiba, Japan) that were co-registered with PET images using PMOD® image analysis software (PMOD Technologies Ltd., Switzerland). The regional radioactivity of each VOI was calculated for each frame and plotted against time. Regional binding potentials relative to non-displaceable radioligands (BP_ND_) of 5-HTRs were estimated using a simplified reference tissue model. Monkeys were scanned with and without drug-treatment condition on different days.

Parametric images of the BP_ND_ were constructed using the original multilinear reference tissue model (48). Individual structural MR images were registered to the Yerkes19 macaque template (49, 50) using FMRIB’ s linear registration tool (FLIRT) and FMRIB’ s nonlinear registration tool (FNIRT) implemented in FSL software (FMRIB’ s Software Library, http://www.fmrib.ox.ac.uk/fsl) (51). BP_ND_ images were then normalized to the template using structural MRI-to-template matrices. Average BP maps across monkeys were plotted on the surface map (52) using the Connectome Workbench (53).

Occupancy levels were determined from the degree of BP_ND_ reduction by antagonists (54). 5-HT receptor occupancy was estimated as follows:

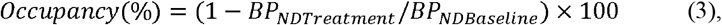

where BP_ND Baseline_ and BP_ND Treatment_ are BP_ND_ measured without (baseline) and with an antagonist, respectively. The target VOI was the visual cortex for 5-HT_1B_R and the global cortex for other subtypes.

### Behavioral tasks and testing procedures

A total of eight monkeys were used for the behavioral study (see Table S1). For all behavioral training and testing, each monkey sat in a primate chair inside a sound-attenuated dark room. Visual stimuli were presented on a computer video monitor in front of the monkey. Behavioral control and data acquisition were performed using a QNX-based Real-time Experimentation data acquisition system (REX; Laboratory of Sensorimotor Research, National Eye Institute) and commercially available software (Presentation, Neurobehavioral Systems). We used three types of behavioral tasks, *reward-size* and *work/delay*, as previously described (2, 55). Both tasks consisted of color discrimination trials (see Figs 2A and 4A). Each trial began when the monkey touched a bar mounted at the front of the chair. The monkey was required to release the bar between 200 and 1,000 ms after a red spot (wait signal) turned green (go signal). Following correctly performed trials, the spot then turned blue (correct signal). A visual cue was presented at the beginning of each color discrimination trial (500 ms before the red spot appeared).

In the reward-size task, a reward of 1, 2, 4, or 8 drops of water (1 drop = ∼0.1 mL) was delivered immediately after the blue signal. Each reward size was selected randomly with equal probability. The visual cue presented at the beginning of the trial indicated the number of drops for the reward (Fig 2A).

In the work/delay task, a water reward (∼0.25 mL) was delivered immediately after each correct signal, after an additional one or two instrumental trials (work trial), or after a delay period (delay trials). The visual cue indicated both the trial type and cost to obtain a reward (Fig 5A). Pattern cues indicated delay trials with the timing of reward delivery after a correct performance: immediately (0.3 s, 0.2–0.4 s; mean, range), after a short delay (3.6 s, 3.0–4.2 s), or after a long delay (7.2 s, 6.0– 8.4 s). Grayscale cues indicated work trials with the number of trials the monkey would have to perform to obtain a reward. We set delay durations to be equivalent to the duration for one or two trials of color discrimination so that we could directly compare the cost of one or two arbitrary units (cost unit; CU).

If the monkey released the bar before the green target appeared or within 200 ms after the green target appeared or failed to respond within 1 s after the green target appeared, we regarded the trial as a “refusal trial”. At this point, all visual stimuli disappeared, the trial was terminated immediately, and after a 1-s inter-trial interval, the trial was repeated. Our behavioral measurement for the motivational value of outcome was the proportion of refusal trials. Before each testing session, monkeys were subjected to ∼22 hours of water restriction in their home cage. Each session continued until the monkey would no longer initiate a new trial (usually less than 100 min). Before this experiment, all monkeys were trained to perform color discrimination trials in the cued multi-trial reward schedule task for more than 3 months. The monkeys were tested with the work/delay task for one or two daily sessions as training to become familiar with the cueing condition. Each monkey was tested from Monday to Friday. Treatment with 5-HTR antagonist or saline (as a control) were performed one day per week.

### Behavioral data analysis

All data and statistical analyses were performed using the R statistical computing environment (R Development Core Team, 2004). The average error rate for each trial type was calculated for each daily session, with error rates in each trial type being defined as the number of error trials divided by the total number of trials of that given type. The monkeys sometimes made many errors at the beginning of the daily session, probably due to high motivation/impatience; therefore, we excluded data until the first successful trial in these cases. A trial was considered an error trial if the monkey released the bar either before or within 200 ms after the appearance of the green target (early release) or failed to respond within 1 s after the green target (late release). We did not distinguish between the two types of errors and used their sum except for the error pattern analysis. We performed repeated-measures ANOVAs to test the effect of treatment × reward size (for data in the reward-size task) or treatment × cost type × remaining cost (for data in the work/delay task) on RT and error pattern. Post hoc comparisons were performed using Tukey HSD test, and a priori statistical significance was set at *p* = 0.05.

We used refusal rates to estimate the level of motivation because the refusal rates of these tasks (*E*) are inversely related to the value for action (2). In the reward-size task, we used the inverse function (Eq. 1). We fit the data to LMMs (56), in which the random effects across 5-HT depletion conditions (i.e., cCPA-day1 and -day2) on parameter *a* and/or intercept *e* (Fig 1) were nested. Model selection was based on Akaike’ s information criterion (AIC), an estimator of in-sample prediction error for nested models (Table S2). Using the selected model, parameter *a* and intercept *e* were estimated individually. Parameter *a* was then normalized to the value in the nontreated condition (control, CON). LMMs were also applied for correlation analysis between refusal rate and RT (Fig 2D), where four statistical models were nested to account for the presence or absence of random effects of subjects and treatment conditions and the best-fit model was selected based on AIC (Table S3).

To examine the effects of satiation, each session was divided into three based on normalized cumulative reward, *R*_*cum*_. Mean error rates in the reward size task across 11 sessions were then fit to the following model:

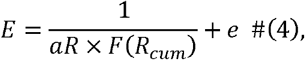

where the satiation effect, *F(R*_*cum*_), as the reward value was exponentially decaying in *R*_*cum*_ at a constant λ (55):

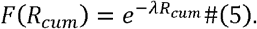

To estimate the effect of 5-HTR blockade on parameters, we also used LMMs. Models were nested to account for the presence or absence of random effects, random effects of treatment conditions, and subjects (see Table S4). The best model was selected based on the AIC for the entire dataset, which is the sum of the results for the regression of each unit nested by individual and/or treatment condition.

In the work/delay task, we used linear models to estimate the effect of remaining cost, i.e., workloads and delay, as previously described (15, 17):

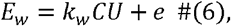

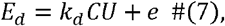

where *E*_*w*_ and *E*_*d*_ are the error rates, and *k*_*w*_ and *k*_*d*_ are cost factors for work and delay trials, respectively. *CU* is the number of remaining cost units, and *e* is the intercept. We simultaneously fitted a pair of these linear models to the data by sum-of-squares minimization without weighting.

We also used LMMs to estimate the effect of 5-HTR blockade on discounting parameters. We imposed the constraint that the intercept (*e*) has the same value between cost types and assumed a basic statistical model in which the random effects of blocking condition independently affect the regression coefficients. Other methods were the same as used for data in the reward size task. Models and results are reported in Table S6.

## Data availability

The data underling all the figures will be available on the following public repository: http://github.com/minamimoto-lab/2023-Yukiko-5HTR.

## Acknowledgements

We thank R. Suma, T. Sugii, R. Yamaguchi, Y. Matsuda, and J. Kamei for their technical assistance. We also thank S. Bouret for his comments on the earlier version of the manuscript. This work was supported by JSPS/MEXT KAKENHI Grant Numbers JP22K07339 (to Yukiko Hori) and JP26120733, JP18H04037 and JP20H05955 (to T.M.). The two Japanese monkeys used in this study were provided by National Bio-Resource Project “Japanese Monkeys” of MEXT, Japan.

