## Supplementary material for "Reduced serotonergic transmission alters sensitivity to cost and reward via 5-HT_1A_ and 5-HT_1B_ receptors in monkeys": Figs S1-S6 and Tables S1-S5

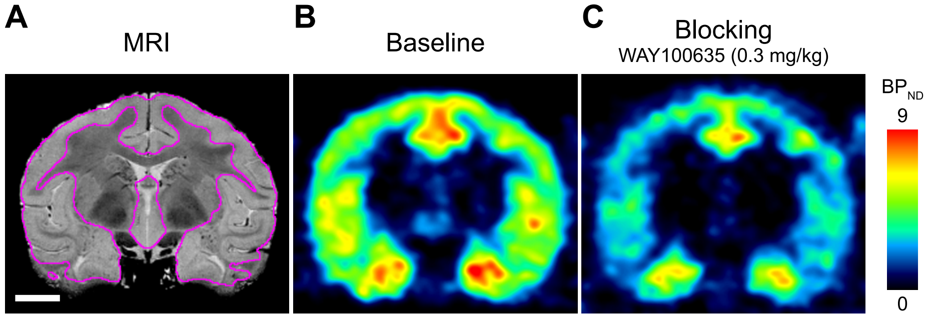


**Fig S1. Occupancy estimation.** Example of occupancy estimation based on the measurement of reduction in specific tracer binding. A. Representative coronal section of an MR image showing the region of interest (ROI, marked by the purple line) for the binding measurement. B and C. Representative coronal sections of parametric PET image showing the specific binding ($BP_{ND}$) of [$^{11}$C]WAY100635 obtained from monkey CH under baseline (B) and the blocking condition with cold WAY100635 (0.3 mg/kg, i.m.) (C). Occupancy was determined as the ratio of reduced specific binding to baseline [i.e., ($BP_{ND}^{baseline}$ – $BP_{ND}^{block}$)/ $BP_{ND}^{baseline}$]. In this case, the reduction in specific binding was 36.7%.

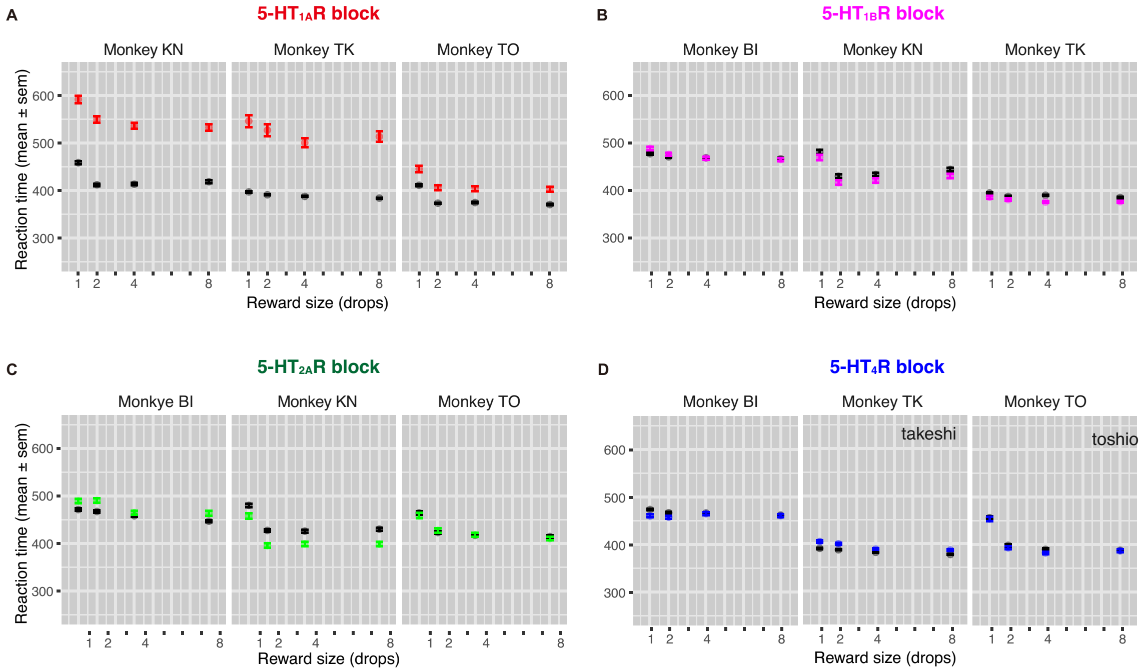


**Fig S2. Effect of 5-HTR blockade on reaction time in reward-size task.** Mean reaction time as function of reward size for control (black) and 5-HTR blockade conditions (color). (A) 5-$HT_{1A}R$ blockade, (B) 5-$HT_{1B}R$ blockade, (C) 5-$HT_{2A}R$ blockade, and (D) 5-$HT_4R$ blockade.

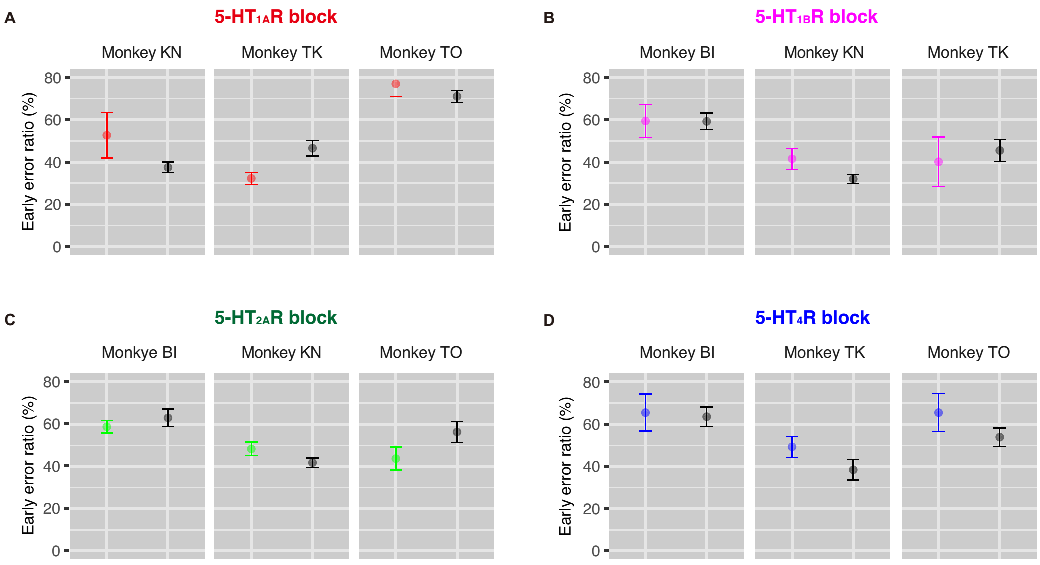


**Fig S3. Effect of 5-HTR blockade on error pattern in reward-size task.**

Early release rate (mean ± SEM) as function of reward size for control (black) and 5-HTR blockade conditions (color). (A) 5-$HT_{1A}$R blockade, (B) 5-$HT_{1B}$R blockade, (C) 5-$HT_{2A}$R blockade, and (D) 5-$HT_4$R blockade.

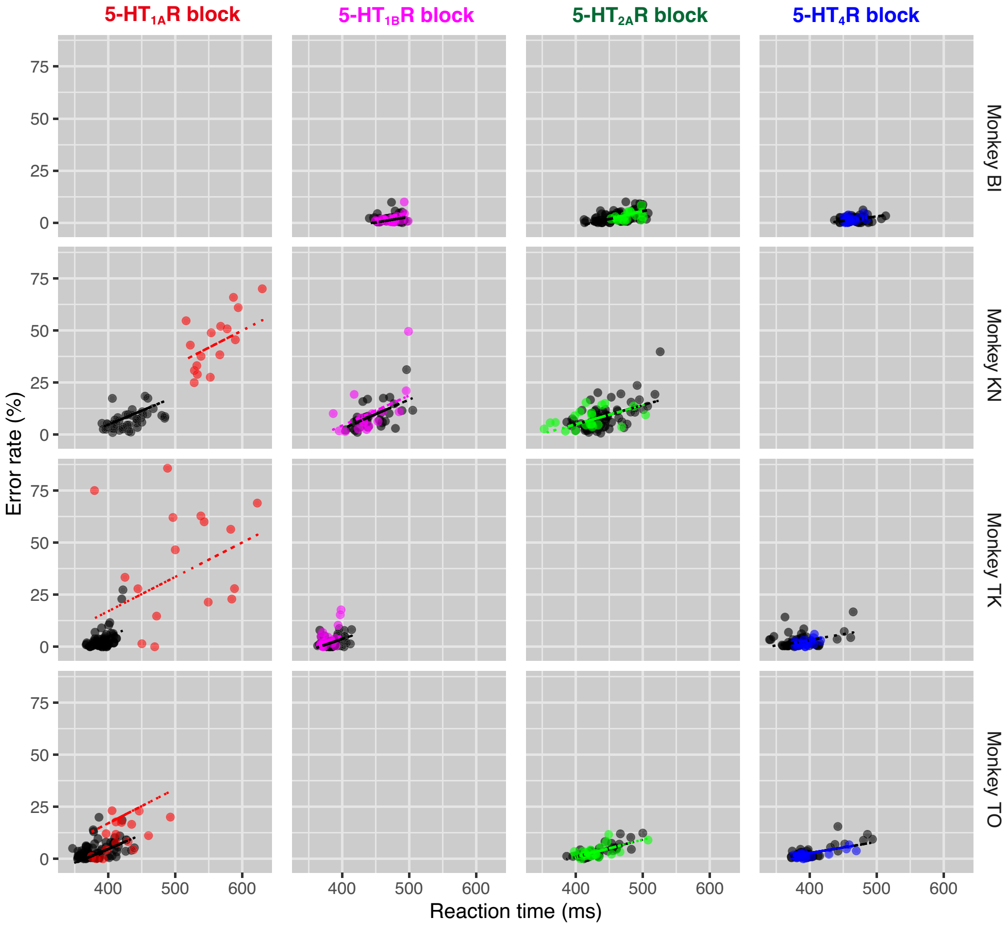


**Fig S4. Effect of 5-HTR blockade on the relationship between refusal rate and reaction time in the reward-size task.** Relationship between refusal rate and mean reaction time for each reward size in session-by-session under control and 5-HTR blockade conditions for each monkey. Colors indicate treatment condition. Lines represent best-fit linear regression models to explain the data (Table S5).

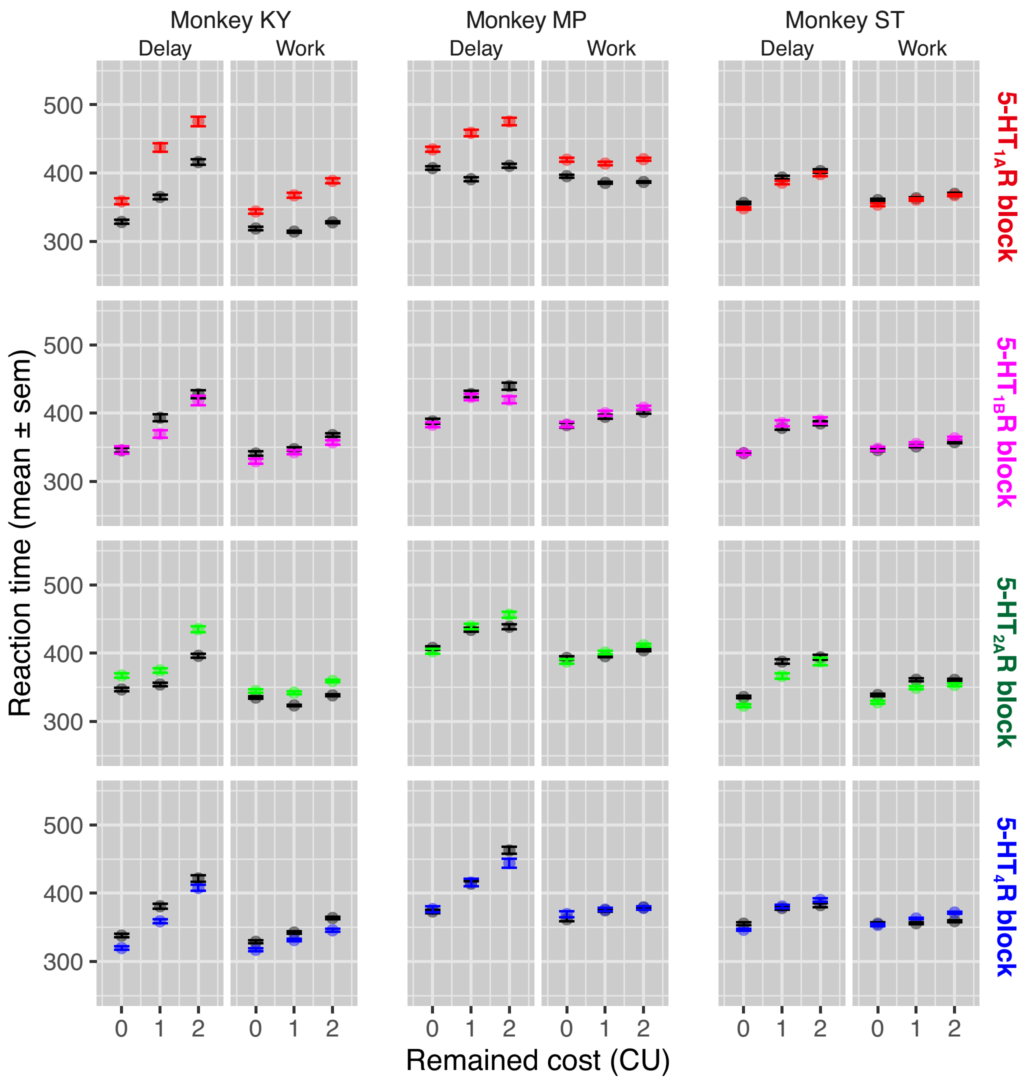


**Fig S5. Effect of 5-HTR blockade on reaction time in delay/workload task.** Mean reaction time as function of reward size for control (black) and 5-HTR blockade conditions (color) in delay and work trials. (A) 5-$HT_{1A}$ blockade, (B) 5-$HT_{1B}$R blockade, (C) 5-HT2AR blockade, and (D) 5-$HT_4$R blockade.

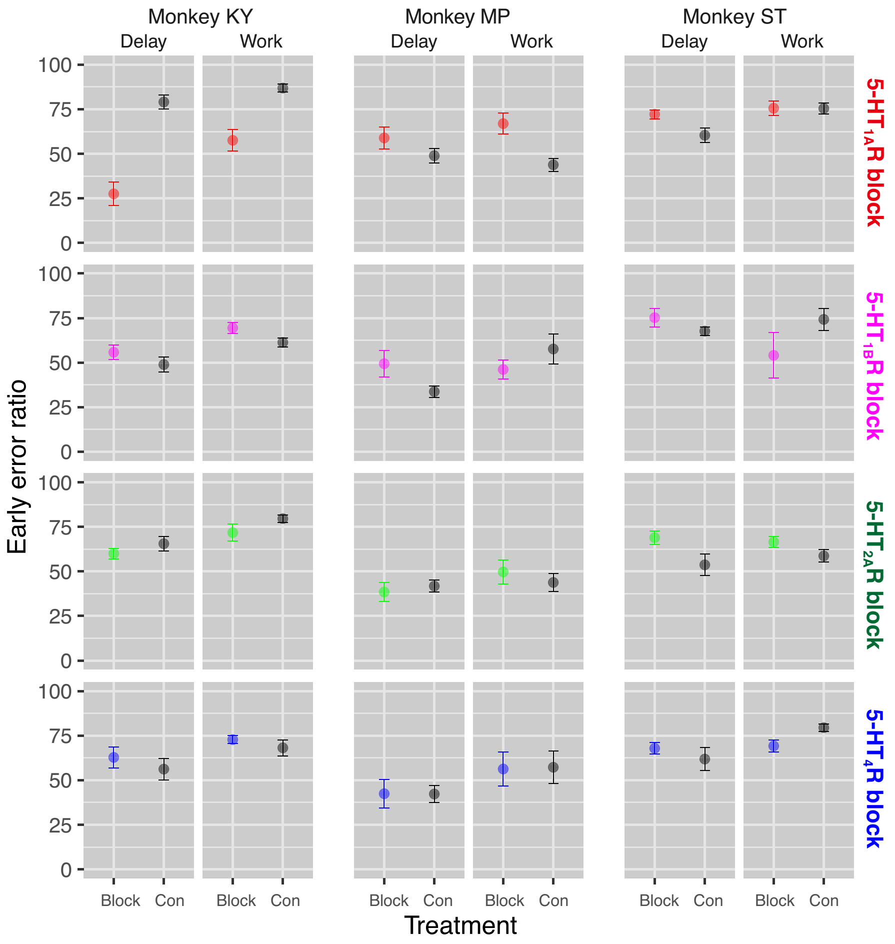


**Fig S6. Effect of 5-HTR blockade on error pattern in work/delay task.**

Late release rate (mean ± SEM) as function of reward size for control (black) and 5-HTR blockade conditions (color). (A) 5-$HT_{1A}$R blockade, (B) 5-$HT_{1B}$R blockade, (C) 5-$HT_{2A}$R blockade, and (D) 5-$HT_4$R blockade.

**Supplementary information**

**Table S1. Summary of subjects used in this study**

| ID | Species | Sex | BW (kg) | Age (y) | 5-HT depletion | | 5-HTR blockade | | |
|---|---|---|---|---|---|---|---|---|---|
| | | | | | CSF | RS task | PET | RS task | W/D task |
| BI | R | M | 7.9 | 7 | | | | ✔ | |
| BO | R | M | 5.7 | 7 | | ✔ | | ✔ | |
| CH | R | M | 7.3 | 4 | | | ✔ | | |
| DR | R | M | 5.7 | 5 | | | ✔ | | |
| JA | R | M | 8.5 | 9 | | | ✔ | | |
| KN | R | M | 6.1 | 12 | | ✔ | | | |
| KY | R | M | 5.9 | 11 | | ✔ | | | ✔ |
| MP | J | M | 7.7 | 7 | | | | | ✔ |
| PE | R | M | 4.5 | 6 | ✔ | | | | |
| SA | R | M | 6.1 | 5 | ✔ | | ✔ | | |
| ST | R | M | 6.2 | 12 | | ✔ | ✔ | | ✔ |
| TA | R | M | 7.7 | 9 | | | ✔ | ✔ | |
| TN | R | M | 6.7 | 7 | | | ✔ | | |
| TO | R | M | 7.0 | 9 | | | | ✔ | |
| TS | R | M | 5.1 | 8 | | | ✔ | | |
| PE | J | M | 7.5 | 5 | | | ✔ | | |
| Total | 16 | | | | 2 | 4 | 9 | 4 | 3 |

Checkmarks indicate the subject was used the experiment. ID, monkey; Species (J, Japanese; R, Resus); Sex (F, Female; M, Male); BW, body weight; CSF, cerebrospinal fluid; RS task, reward-size task; W/D task, work/delay task.

**Table S2. Model comparison for the effect of 5-HT depletion on refusal rate (for Fig 1E)**

| | Model | AIC | ΔAIC |
|---|---|---|---|
| #1 | *E = 1/a(cond) R* | 27.8 | 14.8 |
| #2 | *E = 1/a(cond) R + e* | 23.2 | 10.2 |
| #3 | *E = 1/a(cond) R + e(cond)* | 15.7 | 2.7 |
| **#4** | ***E = 1/a(cond) R + e(cond)**** | **13.0** | **0** |
| #5 | *E = 1/a R + e(cond)* | 20.2 | 6.8 |

*a*(*cond*) and *e*(*cond*) indicate the random effects of 5-HT depletion conditions on parameters *a* and *e*, respectively. The random effect of Model #4 was assumed to be a two-dimensional normal distribution on *a* and *e*. AIC (Akaike's Information Criterion) is a relative measure of the quality for Models #1–#5. ΔAIC denotes the difference from the minimum AIC.

**Table S3. Model comparison for the effect of 5-HT depletion on the relationship between refusal rate and RT in reward-size task (for Fig 2D)**

| | Model | AIC | ΔAIC |
|---|---|---|---|
| #1 | *E ~ RT* | -125 | 4 |
| #2 | *E ~ RT + (RT\|monkey)* | -114 | 15 |
| #3 | *E ~ RT + (RT\|cond)* | -118 | 11 |
| **#4** | ***E ~ RT + (RT\|monkey) + (RT\|cond)*** | **-129** | **0** |

*(RT|*)* indicates random effects on regression parameters. *E*, refusal rate; *RT*, reaction time; *cond*, treatment condition (pCPA-day1, pCPA-day2, or control); *monkey*, subject.

**Table S4. Model comparison for the effect of 5-HTR blockade on refusal rate in reward-size task (for Fig 4)**

| | Model | $5\text{-HT}_{1A}$ | | $5\text{-HT}_{1B}$ | | $5\text{-HT}_{2A}$ | | $5\text{-HT}_{4}$ | |
|---|---|---|---|---|---|---|---|---|---|
| | | AIC | ΔAIC | AIC | ΔAIC | AIC | ΔAIC | AIC | ΔAIC |
| #1 | *E* ~ 1/*a R* + *e*(*cond)* | 190.1 | 0.9 | 139.7 | 40.7 | 121.1 | 23.5 | 80.6 | 5.0 |
| #2 | *E* ~ 1/*a R* + *e(cond), with (*1/*a*(*cond*), *e(cond*)) ~ *biNorm* | 194.0 | 4.9 | 145.7 | 46.7 | 127.1 | 29.5 | 84.2 | 8.6 |
| #3 | *E* ~ 1/*a R* + *e*(*cond*), *with* (1/*a*(*monkey*), *e*(*monkey*)) ~ *biNorm* | **189.2** | **0.0** | 106.0 | 7.0 | 99.5 | 2.0 | 80.2 | 4.5 |
| #4 | *E* ~ 1/*a R* + *e*(*cond*), *with* (1/*a*\|(*cond, monkey*)) ~ *biNorm* | 191.5 | 2.3 | 105.0 | 6.0 | 105.5 | 8.0 | 80.7 | 5.0 |
| #5 | *E* ~ 1/*a*(*cond*) *R* + *e* | 191.7 | 2.5 | 139.7 | 40.7 | 121.1 | 23.5 | 78.2 | 2.6 |
| #6 | *E* ~ 1/*a*(*cond*) *R* + *e* , *with* (1/*a*(*monkey*), *e*(*monkey*)) ~ *biNorm* | 191.9 | 2.7 | **99.0** | **0.0** | 99.5 | 2.0 | **75.6** | **0.0** |
| #7 | *E* ~ 1/*a*(*monkey*) *R* + *e* | 203.8 | 14.6 | 104.7 | 5.7 | 98.7 | 1.2 | 76.0 | 0.3 |
| #8 | *E* ~ 1/*a*(*cond*) *R* + *e*(*cond*), *with* (1/*a*(*cond*), *e*(*cond*)) ~ *biNorm* | 192.0 | 2.9 | 143.7 | 44.7 | 125.1 | 27.5 | 82.2 | 6.6 |
| #9 | *E* ~ 1/*a*(*cond*) *R* + *e*(*cond*), *with* (1/*a*¥(*cond, monkey*), *e*\|(*cond, monkey*)) ~ *biNorm* | 189.5 | 0.3 | 103.0 | 4.0 | 103.5 | 6.0 | 78.7 | 3.0 |
| #10 | *E* ~ 1/*a*(*monkey*) *R* + *e*(*monkey*), *with* (1/*a, e*) ~ *biNorm* | 207.5 | 18.3 | 106.3 | 7.3 | **97.5** | **0.0** | 79.7 | 4.1 |
| #11 | *E* ~ 1/*a R* + *e*(*monkey*) | 203.7 | 14.5 | 122.2 | 23.2 | 101.7 | 4.1 | 79.9 | 4.3 |

*a(cond)* and *e(cond)* indicate the random effects of 5-HTR blocking conditions on parameters *a* and *e*, respectively. AIC (Akaike's Information Criterion) is a relative measure of the quality of Models #1–#11. Δ AIC denotes the difference from the minimum AIC.

**Table S5. Model comparison for the effect of 5-HTR blockade on the relationship between refusal rate and RT in reward-size task (for Fig. S4)**

| | | $5\text{-HT}_{1A}$ | | $5\text{-HT}_{1B}$ | | $5\text{-HT}_{2A}$ | | $5\text{-HT}_{4}$ | |
|---|---|---|---|---|---|---|---|---|---|
| | Model | AIC | ΔAIC | AIC | ΔAIC | AIC | ΔAIC | AIC | ΔAIC |
| #1 | $E + E_0$ | 1886 | 237 | 1015 | 70 | 1361 | 134 | 873 | 40 |
| #2 | $E \sim RT + E_0$ | 1675 | 26 | 1007 | 62 | 1340 | 114 | 874 | 41 |
| #3 | *E ~ RT + E0 + E0\|monkey* | 1675 | 26 | 948 | 3 | 1228 | 2 | 834 | 1 |
| #4 | *E ~ RT + E0 + E0\|RT + RT\|monkey* | 1678 | 29 | 946 | 1 | **1226** | **0** | **833** | **0** |
| #5 | *E ~ RT + E0 + E0\|monkey* | 1677 | 28 | 948 | 3 | 1230 | 3 | 835 | 2 |
| #6 | *E ~ RT + E0 + E0\|cond* | 1651 | 2 | 1019 | 74 | 1348 | 122 | 890 | 58 |
| #7 | *E ~ RT + E0 + E0\|RT + RT\|cond* | **1649** | **0** | 1015 | 70 | 1350 | 123 | 887 | 54 |
| #8 | *E ~ RT + E0 + E0\|cond* | 1655 | 6 | 1015 | 70 | 1350 | 123 | 887 | 54 |
| #9 | *E ~ RT \|monkey+ E0 + E0\|cond)* | 1663 | 15 | 949 | 4 | 1246 | 20 | 836 | 3 |
| #10 | *E ~ RT\|monkey + E0 + E0\|(monkey*cond)* | 1657 | 9 | **945** | **0** | 1247 | 21 | 833 | 0 |
| #11 | *E ~ RT\|monkey + E0* | 1655 | 6 | 947 | 2 | 1230 | 4 | 836 | 3 |
| #12 | *E ~ RT\|monkey + E0 + E0 (RT\|cond)* | 1652 | 3 | 964 | 18 | 1247 | 21 | 852 | 19 |
| #13 | $E \sim RT\|(monkey\|cond) + E_0$ | 1651 | 2 | 959 | 14 | 1245 | 19 | 849 | 16 |
| #14 | $E \sim RT\|monkey + E_0 + E_0\|monkey + E_0\|cond$ | 1656 | 7 | 947 | 2 | 1233 | 7 | 836 | 3 |
| #15 | $E \sim RT\|cond + E_0 + E_0\|cond + E_0\|monkey$ | 1653 | 4 | 951 | 6 | 1232 | 6 | 838 | 5 |
| #16 | $E \sim RT\|cond + E_0 + E_0\|(cond * monkey)$ | 1651 | 2 | 947 | 2 | 1232 | 5 | 836 | 3 |
| #17 | $E \sim RT\|cond + E_0 + RT\|monkey$ | 1656 | 7 | 947 | 2 | 1232 | 5 | 836 | 3 |

*CU* and $E_0$ indicate the remaining cost and intercept, respectively. *a(cond)* and *e(cond)* indicate the random effects of 5-HTR blocking conditions on parameters *a* and *e*, respectively. AIC (Akaike's Information Criterion) is a relative measure of the quality of the models. Δ AIC denotes the difference from the minimum AIC
